# Genome-wide analysis in UK Biobank identifies four loci associated with mood instability and genetic correlation with major depressive disorder, anxiety disorder and schizophrenia

**DOI:** 10.1101/117796

**Authors:** Joey Ward, Rona J. Strawbridge, Mark E. S. Bailey, Nicholas Graham, Ferguson Amy, Donald M. Lyall, Breda Cullen DClinPsy, Laura M. Pidgeon, Jonathan Cavanagh, Daniel F. Mackay, Jill P. Pell, Michael O’Donovan, Valentina Escott-Price, Daniel J. Smith

## Abstract

Mood instability is a core clinical feature of affective and psychotic disorders. In keeping with the Research Domain Criteria (RDoC) approach, it may be a useful construct for identifying biology that cuts across psychiatric categories. We aimed to investigate the biological validity of a simple measure of mood instability and evaluate its genetic relationship with several psychiatric disorders, including major depressive disorder (MDD), bipolar disorder (BD), schizophrenia, attention deficit hyperactivity disorder (ADHD), anxiety disorder and post-traumatic stress disorder (PTSD). We conducted a genome-wide association study (GWAS) of mood instability in 53,525 cases and 60,443 controls from UK Biobank, identifying four independently-associated loci (on chromosomes eight, nine, 14 and 18), and a common single nucleotide polymorphism (SNP)-based heritability estimate of approximately 8%. We found a strong genetic correlation between mood instability and MDD (r_g_=0.60, SE=0.07, p=8.95 × 10^−17^) and a small but significant genetic correlation with both schizophrenia (r_g_=0.11, SE=0.04, p=0.01) and anxiety disorders (r_g_=0.28, SE=0.14, p=0.04), although no genetic correlation with BD, ADHD or PTSD. Several genes at the associated loci may have a role in mood instability, including the *DCC netrin 1 receptor (DCC*) gene, *eukaryotic translation initiation factor 2B subunit beta (eIF2B2*), *placental growth factor (PGF*), and *protein tyrosine phosphatase, receptor type D (PTPRD*). Strengths of this study include the very large sample size, but our measure of mood instability may be limited by the use of a single question. Overall, this work suggests a polygenic basis for mood instability. This simple measure can be obtained in very large samples; our findings suggest that doing so may offer the opportunity to illuminate the fundamental biology of mood regulation.

## Introduction

Mood instability is a common clinical feature of affective and psychotic disorders, particularly major depressive disorder (MDD), bipolar disorder (BD) and schizophrenia^1^. It may also be relatively common in the general population, estimated to affect around 13% of individuals^2^. As a dimensional psychopathological trait, it is potentially a useful construct in line with the Research Domain Criteria (RDoC) approach^3^. Mood instability may be of fundamental importance for understanding the pathophysiology of MDD and BD, as well as conditions such as borderline personality disorder, anxiety disorders, attention deficit hyperactivity disorder (ADHD) and psychosis^4^. This trait is reported by 40-60% of individuals with MDD^5^ and is recognised as part of the prodromal stage of BD^6^. In established BD, it is a clinical feature which independently predicts poor functional outcome^7^. Furthermore, general population twin studies suggest that additive genetic effects account for 40% of the variance in measures of affect intensity and 25% of the variance in affective liability^8^.

Population-based studies such as the Adult Psychiatric Morbidity Survey (APMS) have defined mood instability based on responses to a single question, while clinical studies have made use of more detailed rating scales^4^. However, there is a lack of consensus about how best to measure and classify mood instability, and none of the currently available instruments adequately capture intensity, speed and frequency of affective change, or physiological and behavioural correlates. A recent systematic review proposed that mood instability be defined as “rapid oscillations of intense affect, with a difficulty in regulating these oscillations or their behavioural consequences”^9^. Applying this definition will require the future development and validation of a multidimensional assessment of mood instability, which is currently not available.

Within the UK Biobank population cohort of over 0.5 million individuals^10^, the baseline assessment interview contained a question of relevance to mood instability, specifically: *“Does your mood often go up and down?”* This is similar to the question for mood instability used within the APMS (*“Do you have a lot of sudden mood changes, suffered over the last several years”).* Hypothesizing that this simple question taps into pathological mood instability, we predicted it would be more commonly endorsed by individuals within UK Biobank with MDD and BD, compared to individuals with no psychiatric disorder. Moreover, under the hypothesis that this trait has cross-disorder pathophysiological relevance, we predicted that a genome-wide association study (GWAS) might identify shared genetic liability to mood instability and risk for psychiatric disorders in which disordered mood is a feature, including MDD, BD, schizophrenia, ADHD, anxiety disorder and PTSD. Given the size of the sample, we also aimed to identify loci associated with this measure of mood instability.

## Materials and methods

### Sample

UK Biobank is a large cohort of more than 502,000 United Kingdom residents, aged between 40 and 69 years^10^. The aim of UK Biobank is to study the genetic, environmental and lifestyle factors that cause or prevent disease in middle and older age. Baseline assessments occurred over a four-year period, from 2006 to 2010, across 22 United Kingdom (UK) centres. These assessments were comprehensive and included social, cognitive, lifestyle, and physical health measures. For the present study, we used the first genetic data release based on approximately one third of UK Biobank participants. Aiming to maximise homogeneity, we restricted the sample to those who reported being of white UK ancestry (around 95% of the sample).

UK Biobank obtained informed consent from all participants and this study was conducted under generic approval from the NHS National Research Ethics Service (approval letter dated 13 May 2016, Ref 16/NW/0274) and under UK Biobank approvals for application #6553 “Genome-wide association studies of mental health” (PI Daniel Smith).

### Mood instability phenotype

As part of the baseline assessment, UK Biobank participants completed the 12 items of the neuroticism scale from the Eysenck Personality Questionnaire-Revised Short Form (EPQ-R-S)^11^. One of these items assesses mood instability, namely: *“Does your mood often go up and down?”* Participants responding ‘yes’ to this question were considered to be cases of mood instability and those responding ‘no’ were considered controls. From the control sample, we excluded those who reported being on psychotropic medication, and those who reported a physician diagnosis of psychiatric disorder (including MDD, BD, anxiety/panic attacks, ‘nervous breakdown’, schizophrenia and deliberate self-harm/suicide attempt).

After quality control steps (detailed below) and exclusions (3,679 participants responded ‘don’t know’ and 211 responded ‘prefer not to say’), the final sample for genetic analysis comprised 53,525 cases of mood instability and 60,443 controls. Mood instability cases were younger than controls (mean age 55.8 years (SD=8.05) versus 57.7 years (SD=7.74); p<0.0001) and had a greater proportion of females (55.5% versus 49.6%; p<0.0001).

### Genotyping and imputation

In June 2015, UK Biobank released the first set of genotypic data for 152,729 UK Biobank participants. Approximately 67% of this sample was genotyped using the Affymetrix UK Biobank Axiom array (Santa Clara, CA, USA) and the remaining 33% were genotyped using the Affymetrix UK BiLEVE Axiom array. These arrays have over 95% content in common. Only autosomal data were available under the data release. Data were pre-imputed by UK Biobank as fully described in the UK Biobank interim release documentation^12^. Briefly, after removing genotyped SNPs that were outliers, or were multiallelic or of low frequency (minor allele frequency (MAF) <1%), phasing was performed using a modified version of SHAPEIT2 and imputation was carried out using IMPUTE2 algorithms, as implemented in a C++ platform for computational efficiency^13, 14^. Imputation was based upon a merged reference panel of 87,696,888 biallelic variants on 12,570 haplotypes constituted from the 1000 Genomes Phase 3 and UK10K haplotype panels^15^. Variants with MAF <0.001% were excluded from the imputed marker set. Stringent quality control before release was applied by the Wellcome Trust Centre for Human Genetics, as described in UK Biobank documentation^16^.

### Statistical analyses

#### Quality control and association analyses

Before all analyses, further quality control measures were applied. Individuals were removed based on UK Biobank genomic analysis exclusions (Biobank Data Dictionary item #22010), relatedness (#22012: genetic relatedness factor; a random member of each set of individuals with KING-estimated kinship coefficient >0.0442 was removed), gender mismatch (#22001: genetic sex), ancestry (#22006: ethnic grouping; principal component (PC) analysis identified probable Caucasians within those individuals who were self-identified as British and other individuals were removed from the analysis), and quality control failure in the UK BiLEVE study (#22050: UK BiLEVE Affymetrix quality control for samples and #22051: UK BiLEVE genotype quality control for samples). A sample of 113,968 individuals remained for further analyses. Of these, 53,525 were classed as cases and 60,443 were classified as controls. Genotype data were further filtered by removal of SNPs with Hardy–Weinberg equilibrium *P*<10^−6^, with MAF <0.01, with imputation quality score <0.4 and with data on <90% of the sample after excluding genotype calls made with <90% posterior probability, after which 8,797,848 variants were retained.

Association analysis was conducted in PLINK^17^ using logistic regression under a model of additive allelic effects with sex, age, genotyping array and the first 8 PCs (Biobank Data Dictionary items #22009.01 to #22009.08) as covariates. Sex and age were included as covariates because cases and controls differed significantly on these measures. Genetic PCs were included to control for hidden population structure within the sample, and the first 8 PCs, out of 15 available in the Biobank, were selected after visual inspection of each pair of PCs, taking forward only those that resulted in multiple clusters of individuals after excluding individuals self-reporting as being of non-white British ancestry (Biobank Data Dictionary item #22006). Overall, population structure had little impact on mood instability status. The threshold for genome-wide significance was p<5.0 × 10^−8^.

#### Heritability and genetic correlation between mood instability and psychiatric phenotypes

We applied Linkage Disequilibrium Score Regression (LDSR)^18^ to the GWAS summary statistics to estimate SNP heritability (h^2^_SNP_). Genetic correlations between mood instability and MDD, BD, schizophrenia, ADHD, anxiety disorder and PTSD were also evaluated using LDSR^19^ (with unconstrained intercept), a process that corrects for potential sample overlap without relying on the availability of individual genotypes ^18^. For the MDD, BD, schizophrenia, ADHD, anxiety disorder and PTSD phenotypes, we used GWAS summary statistics provided by the Psychiatric Genomics Consortium (http://www.med.unc.edu/pgc/)^20^^-^^25^. Note that for the purposes of these genetic correlation analyses we re-ran the GWAS of mood instability excluding from the cases those 9,865 participants who reported being on psychotropic medication, or who self-reported psychiatric disorder (MDD, BD, anxiety/panic attacks, ‘nervous breakdown’, schizophrenia and deliberate self-harm/suicide attempt). This secondary GWAS output (rather than the primary GWAS reported below) was used for the genetic correlation calculations and for polygenic risk score analyses, the rationale being that this was a more conservative approach which would avoid genetic correlations between mood instability and MDD/BD/schizophrenia/ADHD/anxiety disorders/PTSD being driven by a subset of individuals with psychiatric disorder.

#### Polygenic risk score analysis of MDD, BD and schizophrenia as predictors of mood instability

Polygenic risk scores (PRS) were created using the output of the PCG MDD 29 of 32 cohort GWAS (supplied by the MDD working group of the PGC, http://www.med.unc.edu/pgc/pgc-workgroups), BD GWAS^20^ and schizophrenia GWAS^21^. Five PRS were created for each psychiatric phenotype using p-value cut offs of p<5 × 10^−8^, p<0.01, p<0.05, p<0.1, and p<0.5, with the exception of MDD for which there were no genome wide significant SNPs. Ambiguous SNPs, indels (insertion/deletion mutations) and SNPs with an imputation quality score of less than 0.8 were removed. LD clumping was performed via PLINK on a random sample of 10,000 individuals using an r^2^>0.05 in a 250kb window. SNPs were clumped into sets and filtered, selecting the SNP with the lowest p-value from each set. In the event that 2 or more SNPs from a set had the same p-value, the SNP with the largest beta coefficient was used. PLINK was also used to calculate the PRS to produce a per-allele weighted score with no mean imputation.

#### Polygenic risk score modelling

Only those subjects who were used for the genetic correlations analyses were used in the PRS analyses (that is, PRS analyses also excluded from both case and control groups those individuals in UK Biobank with psychiatric disorder). Modelling was performed in R (version 3.1.2) using the glm function. Full sample and age-stratified analysis models were adjusted for age, sex, chip and PGCs 1- 8, whereas sex-stratified analysis was not adjusted for sex. Scores were then split into deciles using the ntile function of the dplyr package. Model Nagelkerke r^2^ was calculated using the fmsb package.

## Results

### Mood instability in MDD and BD within UK Biobank

In previous work we have identified individuals within UK Biobank with a probable diagnosis of mood disorder, including cases of MDD (sub-divided into single episode MDD, recurrent moderate MDD and recurrent severe MDD) and BD, as well as non-mood disordered controls^26^. These classifications were independent of response to the mood instability question or other questions from the EPQ-R-S. For the group of participants who could be classified in this way, we assessed the proportion with mood instability within each mood disorder category. All mood disorder groups had a significantly greater proportion of individuals with mood instability compared with the control group (Table 1), in which the prevalence was 35.3%. This proportion was highest in the BD group (74.0%) followed by the three MDD groups (71.7% for recurrent severe MDD, 64.2% for recurrent moderate MDD and 43.7% for single episode MDD). There were too few UK Biobank participants with a reliable classification of schizophrenia, ADHD, anxiety disorder or PTSD to allow for an assessment of the prevalence of mood instability in these groups.

**Table 1.** Proportion of individuals with mood instability within mood disorder groups, compared to non-mood disordered controls.

|  | **Mood instability**<br>**N (%)** | **Pearson Chi-**<br>**squared** | **P-value** |
| --- | --- | --- | --- |
| **BD** | 1,180 (74.0) | $1.0 \times 10^3$ | <0.001 |
| **Recurrent MDD, severe** | 6,303 (71.7) | $4.5 \times 10^3$ | <0.001 |
| **Recurrent MDD, moderate** | 9,509 (64.2) | $4.4 \times 10^3$ | <0.001 |
| **Single episode MDD** | 3,403 (43.7) | 221.1 | <0.001 |
| **Non-mood disordered controls** | 30,844 (35.3) | - | - |

*BD bipolar disorder; MDD major depressive disorder*

### GWAS of mood instability

The mood instability GWAS results are summarised in Figure 1 (Manhattan plot), Figure 2 (QQ plot) and Table 2 (genome-wide significant loci associated with mood instability). Regional plots are provided in Figures 3a-3d.

**Figure 1.**
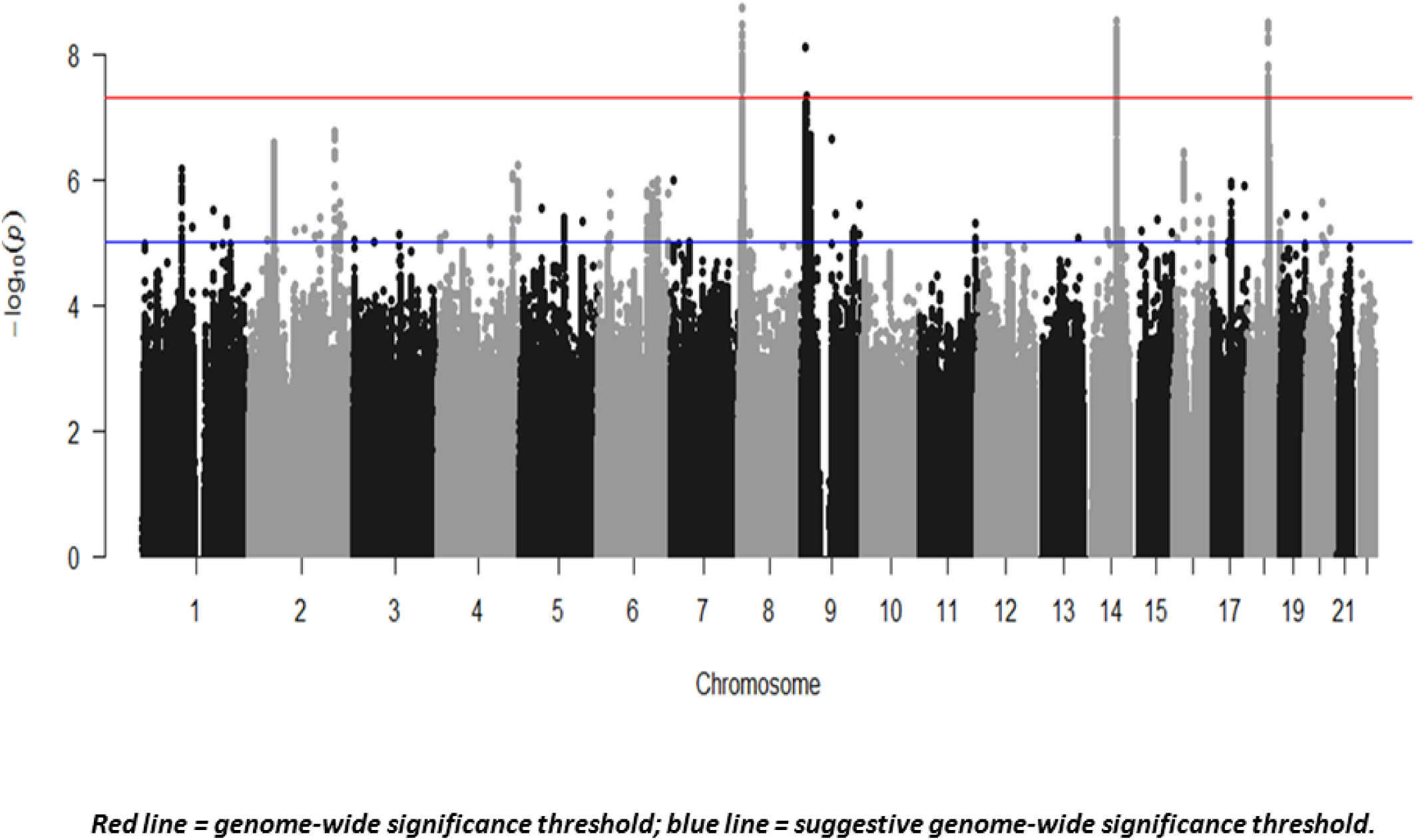
Manhattan plot of GWAS of mood instability in UK Biobank (n=113,968).

**Figure 2.**
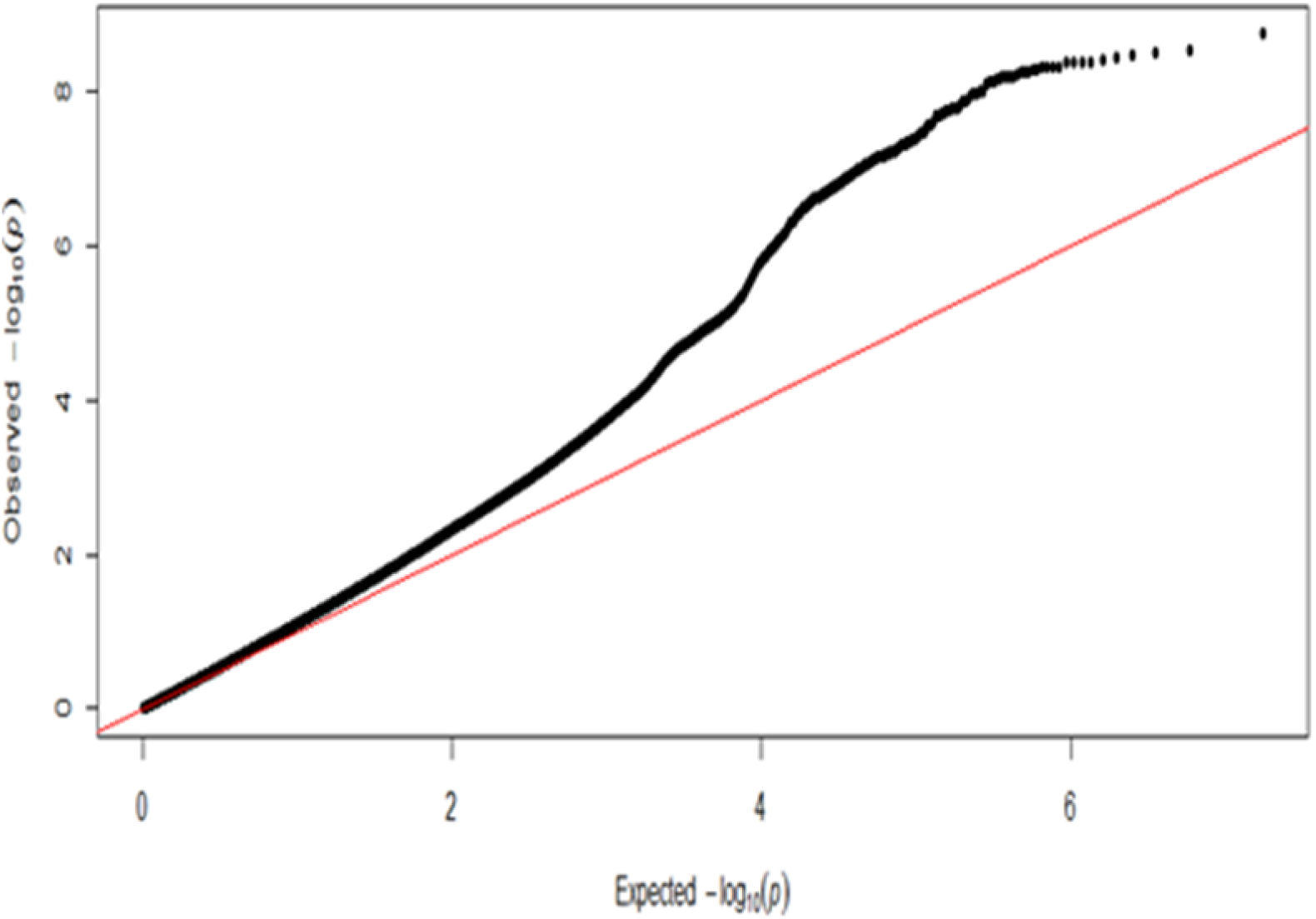
QQ plot for UK Biobank mood instability GWAS results.

**Figures 3a-3d.**
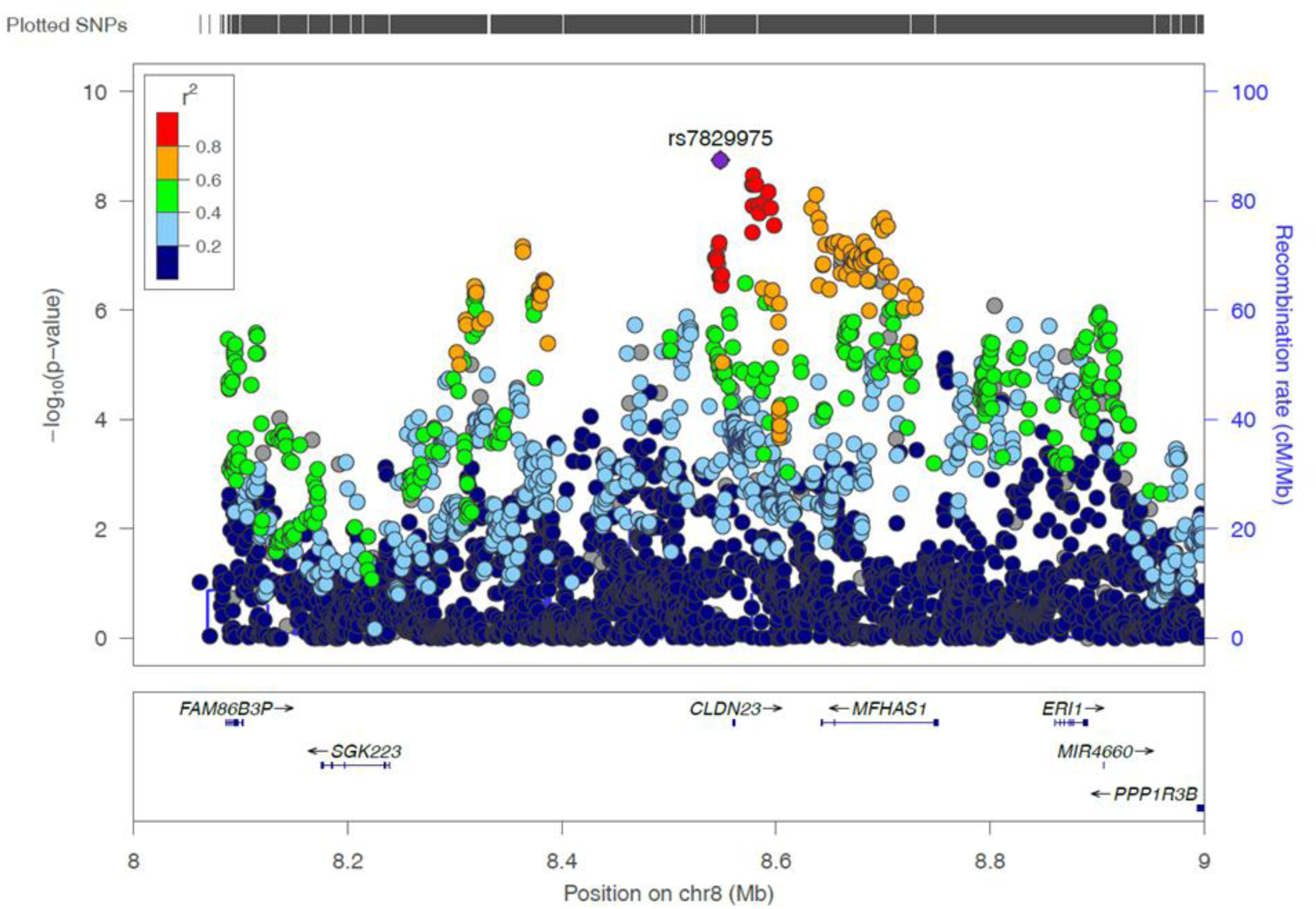
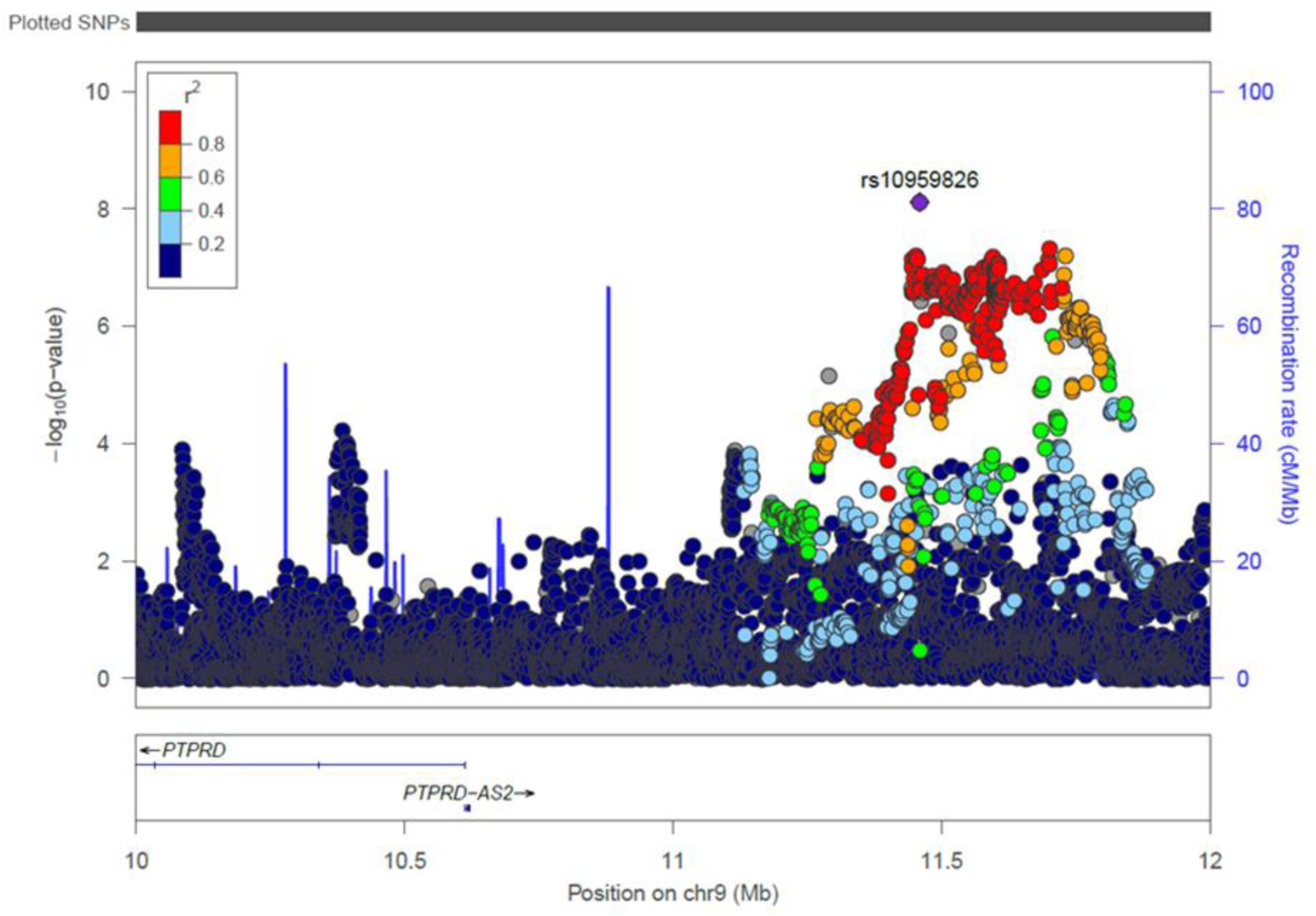
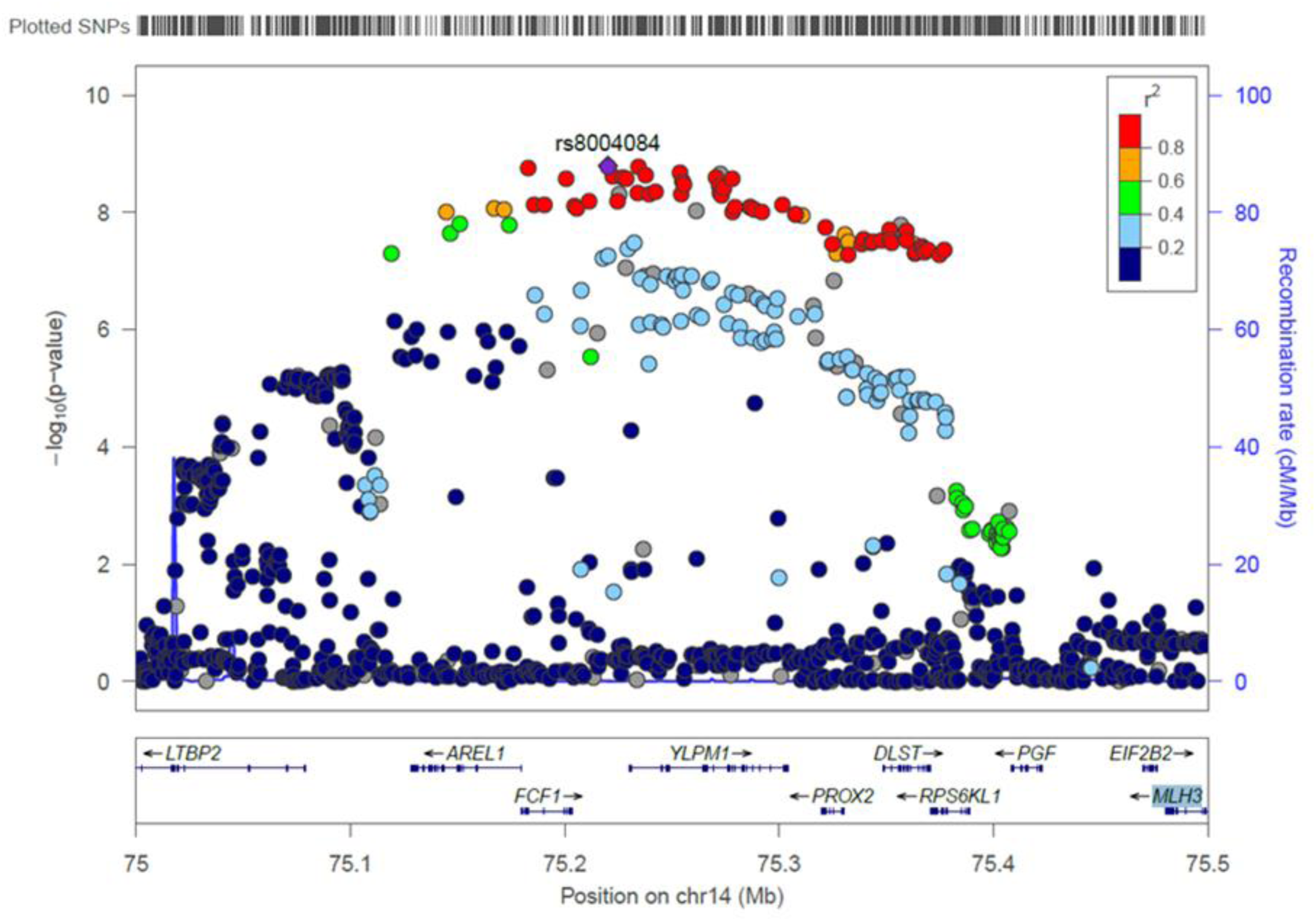
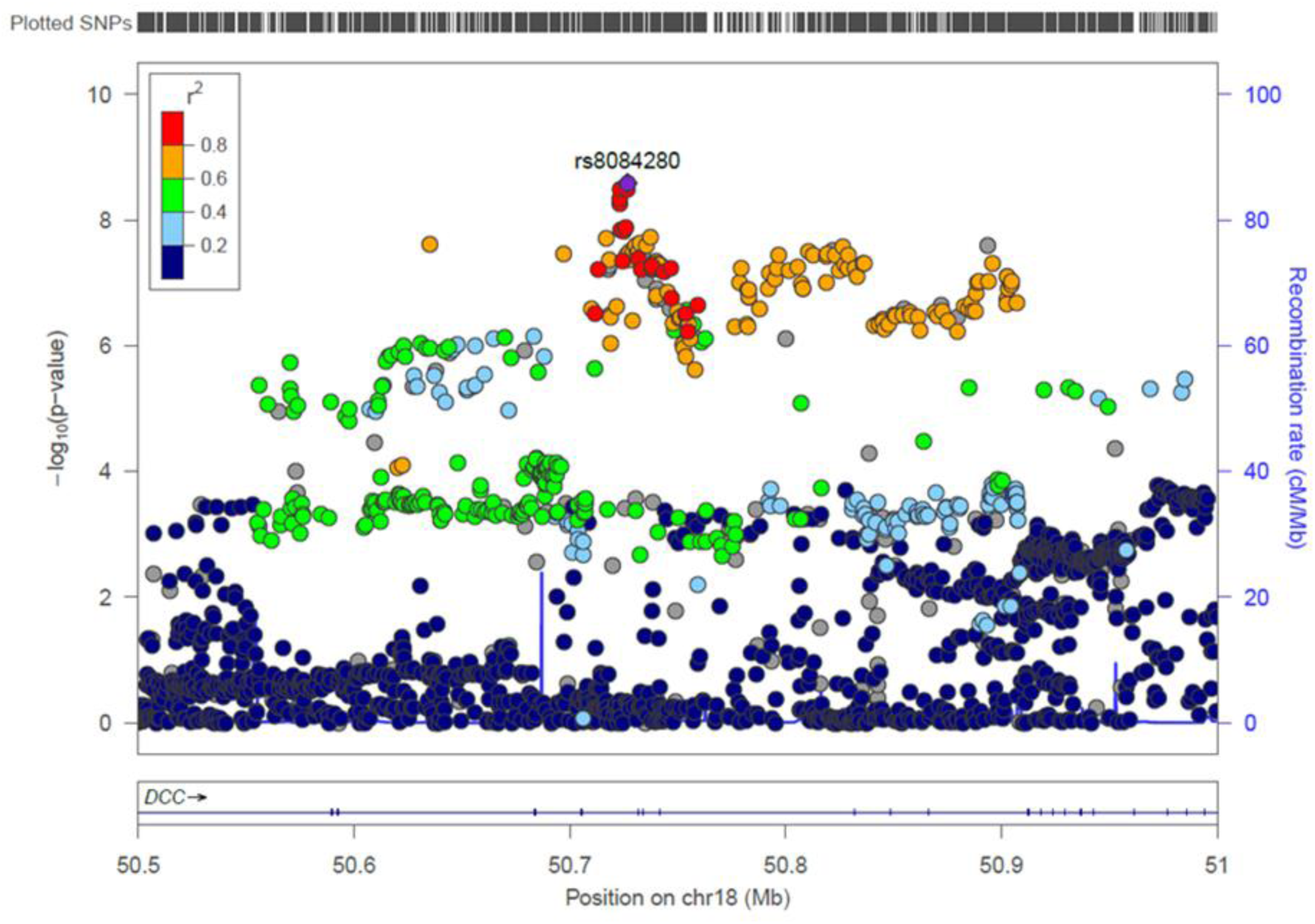
Regional plots of the four genome-wide significant mood instability loci. Figure 3a. Chromosome 8 region 8.5MB-8.8MB Figure 3b. Chromosome 9 region 10MB – 12MB Figure 3c. Chromosome 14 region 75MB-75.5MB Figure 3d. Chromosome 18 region 50.5MB-51MB

**Table 2.** Genome-wide significant loci associated with mood instability in UK Biobank.

| Index SNP | Chr | Position | Risk Allele/Other Allele | RAF | Beta (SE) | P-value | Associated region | Nearby Genes |
| --- | --- | --- | --- | --- | --- | --- | --- | --- |
| rs7829975 | 8 | 8,548,117 | A/T | 0.516 | 0.051 (0.0085) | $1.8 \times 10^{-9}$ | 8,548,117 – 8,704,330 | *CLDN23, MFHAS1* |
| rs10959826 | 9 | 11,459,410 | G/A | 0.785 | 0.060 (0.01) | $7.7 \times 10^{-9}$ | 11,459,410 – 11,701,596 | *PTPRD* |
| rs397852991 | 14 | 75,268,920 | C/CA | 0.606 | 0.053 (0.0088) | $2.98 \times 10^{-9}$ | 75,144,618 – 75,359,229 | *LTBP2, AREL1, FCF1, YLPM1, PROX2, DLST, RPS6KL1, PGF, EIF2B2, MLH3* |
| rs8084280 | 18 | 50,726,749 | T/A | 0.508 | 0.050 (0.0085) | $3.15 \times 10^{-9}$ | 50,635,119 – 50,893,647 | *DCC* |

Shown are LD-independent genome-wide significant SNP associations for mood instability (sorted by genomic position according to NCBI Build 37). Chromosome (Chr) and Position denote the location of the index SNP. RAF = risk allele frequency. Beta = logistic regression coefficient for allele1, SE = standard error for Beta. P-value = the probability of getting the derived test statistic under the null hypothesis. The final column indicates protein-coding genes at the associated loci (see regional plots in supplementary information) or, where there are no genes at the associated locus, the nearest gene if less than 1 MB from the locus.

Overall, the GWAS data showed modest deviation in the test statistics compared with the null (*λ*_GC_ =1.13); this was negligible in the context of sample size (*λ*_GC_ 1000=1.002). LDSR suggested that deviation from the null was due to a polygenic architecture in which h^2^_SNP_ accounted for approximately 8% of the population variance in mood instability (observed scale *h*^2^_SNP_=0.077 (SE 0.007)), rather than inflation due to unconstrained population structure (LD regression intercept=0.998 (SE 0.009)).

We observed four independent genomic loci exhibiting genome-wide significant associations with mood instability (Figure 1, Table 2 and Figures 3a-d), on chromosome eight (index SNP rs7829975; *CLDN23 and MFHAS1*), chromosome nine (index SNP rs10959826; *PTPRD*), chromosome 14 (index SNP rs397852991; *LTBP2, AREL1, FCF1, YLPM1, PROX2, DLST, RPS6KL1, PGF, EIF2B2* and *MLH3*) and chromosome 18 (index SNP rs8084280; *DCC*). In total, there were 111 genome-wide significant SNPs across all loci. Given the functional alleles that drive association signals in GWAS may not affect the nearest gene, we use the above gene names to provide a guide to location rather than to imply that altered function or expression of those genes are the sources of the association signals.

We also repeated this GWAS for males and females separately (supplementary material Figure S1 and Figure S2) and for the sample stratified according to median age (age 58 and below, and age 59 and above; supplementary material Figure S3 and Figure S4). No genome-wide significant loci were observed from these stratified analyses, possibly because of reduced power, apart from the retention of a single genome-wide significant finding at rs8084280 on chromosome 18 (the *DCC* gene) for males only (supplementary material Figure S1). There was a high degree of genetic correlation between mood instability in males and females (r_g_=1.02, SE=0.09, p=2.84 × 10^−30^), and between mood instability in the younger and older sub-groups (r_g_=1.02, SE=0.09, p=2.67 × 10^−27^).

Within supplementary materials, we also present the results of the secondary GWAS of mood instability which excluded from the case group 9,865 participants with a psychiatric disorder (supplementary table S1). This GWAS was used to assess for genetic correlation between mood instability and MDD, BD, schizophrenia, ADHD, anxiety disorders and PTSD, and for the polygenic risk score analyses. Supplementary table S1 shows that the risk allele frequencies (RAFs) of the index SNPs within the four genome-wide significant loci from the primary GWAS were very similar to the RAFs for these same SNPs within this secondary GWAS: for rs7829975 the RAF was 0.516 versus 0.523; for rs10959826 it was 0.785 versus 0.789; for rs397852991 it was 0.606 versus 0.673; and for rs8084280 it was 0.508 versus 0.514). However, it should be noted that, perhaps due to a loss of power from excluding 9,865 individuals, only one of these four loci retained genome-wide significance (rs7829975 on chromosome 8).

### Genetic correlation of mood instability with MDD, schizophrenia, BD, ADHD, anxiety disorder and PTSD

We identified strong genetic correlation between mood instability and MDD (rg=0.60, SE=0.07, p=8.95 × 10^−17^) and a smaller, but significant, correlation between mood instability and both schizophrenia (r_g_= 0.11, SE=0.04, p=0.01) and anxiety disorders (r_g_= 0.28, SE=0.14, p=0.04) (Table 3). We did not find significant genetic overlap between mood instability and BD (r_g_=0.01, SE=0.05, p=0.27), ADHD (r_g_= 0.14, SE=0.11, p=0.18) or PTSD (r_g_=0.33, SE=0.17, p=0.06).

**Table 3.** Genetic correlation between mood instability and MDD, schizophrenia, BD, PTSD, ADHD and anxiety disorder.

| Phenotype | R<sub>g</sub> | se | z | p | h<sup>2</sup> obs | h<sup>2</sup> obs se | h<sup>2</sup> int | h<sup>2</sup> int se | Gcov int | Gcov int se |
| --- | --- | --- | --- | --- | --- | --- | --- | --- | --- | --- |
| MDD | 0.6 | 0.07 | 8.32 | 8.95 x10<sup>-17</sup> | 0.11 | 0.01 | 0.99 | 0.008 | -0.0019 | 0.006 |
| Schizophrenia | 0.11 | 0.04 | 2.48 | 0.01 | 0.25 | 0.01 | 1.03 | 0.01 | 0.0008 | 0.007 |
| BD | 0.01 | 0.05 | 0.27 | 0.27 | 0.12 | 0.01 | 1.02 | 0.008 | 0.0069 | 0.005 |
| PTSD | 0.33 | 0.17 | 1.9 | 0.06 | 0.10 | 0.004 | 0.99 | 0.007 | 0.0004 | 0.005 |
| ADHD | 0.14 | 0.11 | 1.35 | 0.18 | 0.4 | 0.15 | 1.01 | 0.01 | 0.0046 | 0.004 |
| Anxiety disorder | 0.28 | 0.14 | 2.04 | 0.04 | 0.06 | 0.03 | 1.00 | 0.01 | 0.01 | 0.005 |

R<sub>g</sub> = genetic correlation with mood instability; SE = standard error of the genetic correlation; Z = the test statistic; P= p-value. h<sup>2</sup> obs = heritability on the observed scale; h<sup>2</sup> obs SE = the standard error of the heritability; h<sup>2</sup> int = intercept of the heritability; h<sup>2</sup> int SE = standard error of the heritability intercept; Gcov int = intercept of the genetic covariance; Gcov int SE = standard error of the genetic covariance intercept. MDD = major depressive disorder; BD = bipolar disorder; PTSD = post-traumatic stress disorder; ADHD = attention deficit hyperactivity disorder.

### Polygenic risk score analysis of MDD, BD and schizophrenia as predictors of mood instability

Using the PRS approach, both MDD and schizophrenia had significant positive correlations with mood instability status (for MDD at p<0.5 PRS threshold: OR=1.029, 95%CI=1.02-1.033, r^2^=0.023, p=1.00 × 10^−34^ and for schizophrenia at p<0.1 PRS threshold: OR=1.009, 95%CI=1.005-1.014, r^2^=0.021, p= 6.71 × 10^−05^) (supplementary material Table S2). There was no evidence of an association between PRS for BD and mood instability. This finding of a positive correlation between PRSs for MDD and schizophrenia and mood instability status (and no such correlation for BD PRS) was consistent across additional analyses stratified for sex and age (supplementary material Tables S3-S6).

## Discussion

We have identified four independent loci associated with mood instability within a large population cohort, in what is to date the only GWAS of this phenotype. We also identified a SNP-based heritability estimate for mood instability of approximately 8%, and a strong genetic correlation between mood instability and MDD, suggesting substantial genetic overlap between mood instability and vulnerability to MDD. There was also a small but significant genetic correlation between mood instability and schizophrenia and between mood instability and anxiety disorders, but no significant genetic correlation with BD, ADHD or PTSD. Polygenic risk score analyses found a positive correlation between genes for both MDD and schizophrenia and mood instability status, but this was not the case for BD.

The strong genetic correlation between mood instability and MDD is of interest because it is consistent with the hypothesis that at least part of the pathophysiology of MDD might include a reduced capacity to effectively regulate affective states. In support of this is evidence that individuals with MDD tend to have maladaptive responses to intense emotions, responding with worry, rumination and self-criticism, which can then exacerbate negative emotional states^27^. This maladaptive pattern of responses is also consistent with our finding of a small but significant genetic correlation between mood instability and both anxiety disorder and schizophrenia.

The lack of genetic correlation between mood instability and BD was unexpected, given that mood instability is considered a core deficit in BD^4^ and was more common in our BD cases than MDD cases. Similarly, a genetic correlation between mood instability, ADHD and PTSD might have been anticipated. This lack of correlation between mood instability and BD/ADHD/PTSD is difficult to account for but might be explained by the relatively under-powered nature of the BD, ADHD and PTSD GWAS analyses, compared to the analyses used for MDD and schizophrenia. It is worth noting that, although not significant, the magnitude of the genetic correlation between mood instability and ADHD was 0.14. Similarly, the genetic correlation between mood instability and PTSD was not significant but had a magnitude of 0.33.

It is well documented that MDD occurs more commonly in females than in males and it is possible that mood instability may be of greater relevance as a cross-cutting phenotype for women compared to men. We therefore carried out a GWAS of mood instability for males and females separately (supplementary material Figure S1 and Figure S2). These stratified analyses found no genome-wide significant loci for females and only one genome-wide significant locus for males (the previously identified locus on chromosome 18). Furthermore, there was perfect genetic correlation between mood instability in males and females. Although these analyses had reduced power, they suggest that there was no evidence for a large number of sex-specific loci for mood instability. Similarly, we carried out GWAS stratified by age, for those in the sample at or below the median age of 58 and for those above age 58 (supplementary material Figure S3 and Figure S4). As with stratification by sex, these age-stratified analyses did not identify any genome-wide significant loci and there was perfect correlation between mood instability in the younger and older sub-groups.

It is not possible to be certain which of the genes within associated loci are likely to be most relevant to the pathophysiology of mood instability but several genes of interest were identified. For example, the lead SNP within the associated region on chromosome 18 lies in intron 9 of the *DCC netrin 1 receptor* (originally named *deleted in colorectal cancer; DCC*) gene, with no other protein-coding genes for >500kb on either side (Figure 3d). *DCC* is the receptor for the guidance cue *netrin-1*, which has a central role in the development of the nervous system, including (but not limited to) the organization and function of mesocorticolimbic dopamine systems^28^. Recent studies have shown a range of human phenotypes associated with loss-of-function mutations in *DCC*, including agenesis of the corpus callosum, learning disabilities and mirror movements, all associated with a large-scale disruption of the development of commissural connectivity and lateralisation^29, 30^. Manitt and colleagues have identified that *DCC* has a role in regulating the connectivity of the medial prefrontal cortex during adolescence and found that *DCC* expression was elevated in the brain tissue of antidepressant-free subjects who committed suicide^31^. This suggests a possible role for *DCC* variants in increasing predisposition to mood instability and mood disorders, as well as related psychopathological phenotypes.

The associated region on chromosome 14 contains at least 10 candidate genes (Table 2 and Figure 3c). One of these is *translation initiation factor 2B subunit beta (EIF2B2*), mutations in which are known to cause a range of clinically heterogeneous leukodystrophies^32^. Reduced white matter integrity has been consistently associated with negative emotionality traits (such as harm avoidance, neuroticism and trait anxiety)^33^, as well as with MDD and BD^34^. It is therefore possible that variation in *EIF2B2* may have a role in mood instability.

Another gene within the associated region on chromosome 14 is *placental growth factor (PGF*), a member of the angiogenic vascular endothelial growth factor (*VEGF*) family^35, 36^, which is expressed at high levels in the placenta and thyroid^37^. *PGF* has a wide range of functions, including embryonic thyroid development^38^ and immune system function^39, 40^, as well as a role in atherosclerosis, angiogenesis in cancer, cutaneous delayed-type hypersensitivity, obesity, rheumatoid arthritis and pre-eclampsia^39,41-44^. *PGF* may be of interest because of the long-established association between thyroid dysfunction and both MDD and BD^45^, along with the recent observation that pre-eclampsia may be a marker for the subsequent development of mood disorders^46^.

Also of interest is the finding that the gene for *protein tyrosine phosphatase, receptor type D (PTPRD*) lies within 1Mb of the associated region on chromosome 9 (Figure 3b). *PTPRD* encodes a receptor-type protein tyrosine phosphatase known to be expressed in brain and with an organising role at a variety of synapses, including those that play a role in synaptic plasticity^47^. As such, it may have a role in a broad range of psychopathology.

Two of the genomic loci associated with mood instability (on chromosomes eight and nine) overlap with loci found to be associated with neuroticism in a recent GWAS and meta-analysis which combined data from the UK Biobank cohort, the Generation Scotland cohort, and a cohort from the Queensland Institute of Medical Research^48^. The neuroticism study made use of scores on the 12- item EPQ-R-S questionnaire, of which one of the questions was the mood instability question used in the present study. This overlap in findings suggests that mood instability is a key component of neuroticism as defined by the EPQ-R-S and that at least some of the gene variants implicated in mood instability are likely to contribute to the broader phenotype of neuroticism. We did not assess for genetic correlation between mood instability and neuroticism using LDSR because both GWAS outputs were predominantly from the same UK Biobank sample.

### Strengths and limitations

To the best of our knowledge, this is the first reported GWAS of mood instability. It has enabled objective estimates of heritability and genetic correlation with important psychiatric disorders to be made for the first time. In the future, genotyping data for the full UK Biobank sample (502,000 participants) will be available. This increased sample size may identify larger estimates of shared variance between mood instability and psychiatric disorders.

Some important limitations of this work are acknowledged. The mood instability phenotype used was based on response to a single-item question (*“Does your mood often go up and down?”*) which may be an imperfect measure of mood instability. Approximately 44% of the whole UK Biobank cohort answered ‘yes’ to this question, a much larger proportion than the 13% of participants classified as having mood instability within the UK APMS^2^. This may be because the assessment of mood instability in the APMS was based on a slightly different question (*“Do you have a lot of sudden mood changes”*) and because respondents had to additionally report that they *“suffered this symptom over the last several years”.* Clearly, a potential limitation of self-report is the possibility of responder bias and, further, a more complete and objectively-assessed measure of mood instability would have been preferable. However, this was not available to us in the UK Biobank phenotype dataset and is unlikely to be feasible to collect within a population cohort of this size.

## Conclusions

Despite a recognition that mood instability is likely to be an important phenotype underpinning a range of psychiatric disorders - particularly mood disorders^4^ - there has to date been very little work on its neural correlates. Early investigations tentatively suggest a role for altered function and/or connectivity of the amygdala^49^ but this is an area which is currently under-developed. It is hoped that our findings will stimulate new research on mood instability, which may be a clinically useful and biologically valid trait that cuts across traditional diagnostic categories^50^.

## Acknowledgements

This research was conducted using the UK Biobank resource. UK Biobank was established by the Wellcome Trust, Medical Research Council, Department of Health, Scottish Government and Northwest Regional Development Agency. UK Biobank has also had funding from the Welsh Assembly Government and the British Heart Foundation. Data collection was funded by UK Biobank. JW is supported by the JMAS Sim Fellowship for depression research from the Royal College of Physicians of Edinburgh (173558). DJS is supported by an Independent Investigator Award from the Brain and Behaviour Research Foundation (21930) and a Lister Prize Fellowship (173096). AF is supported by an MRC Doctoral Training Programme Studentship at the University of Glasgow (MR/K501335/1). The work at Cardiff University was funded by Medical Research Council (MRC) Centre (G0800509) and Program Grants (G0801418). The funders had no role in the design or analysis of this study, decision to publish, or preparation of the manuscript.

## Figure legends

**Figure S1.**
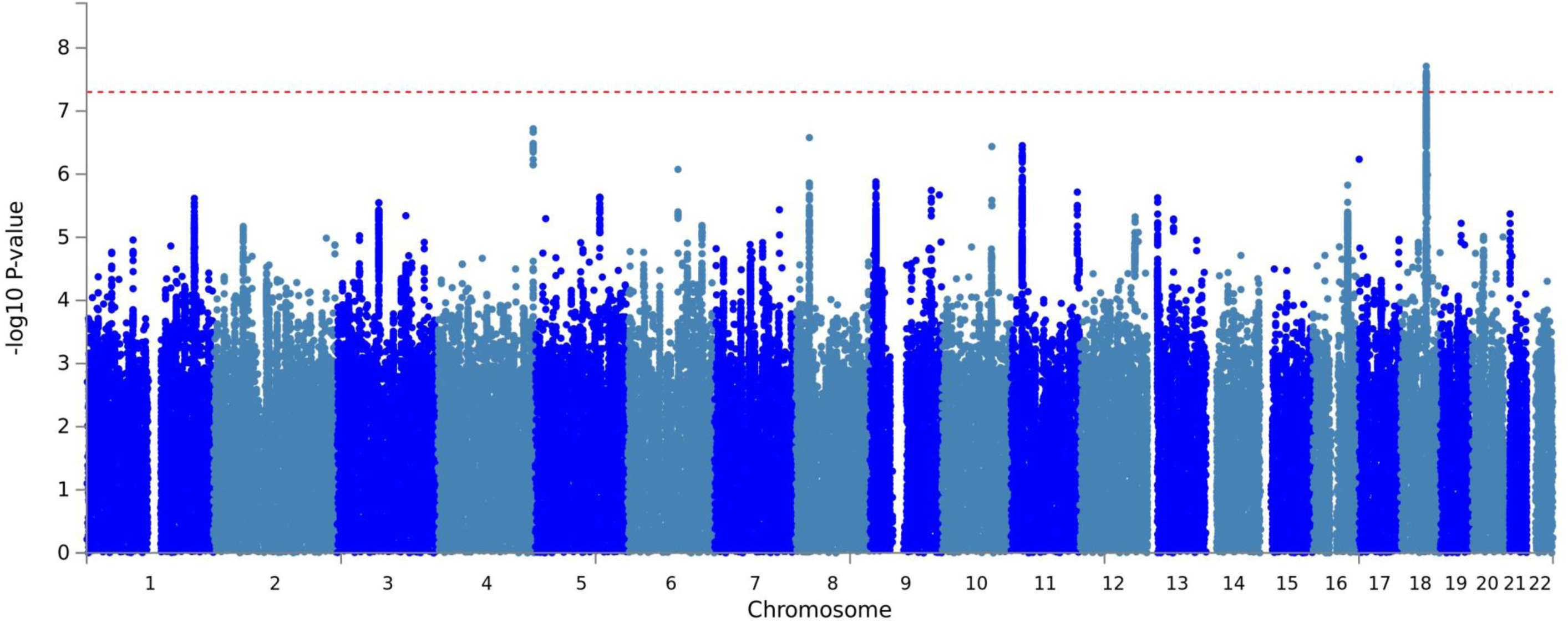
Manhattan plot of GWAS of mood instability in UK Biobank (males only).

**Figure S2.**
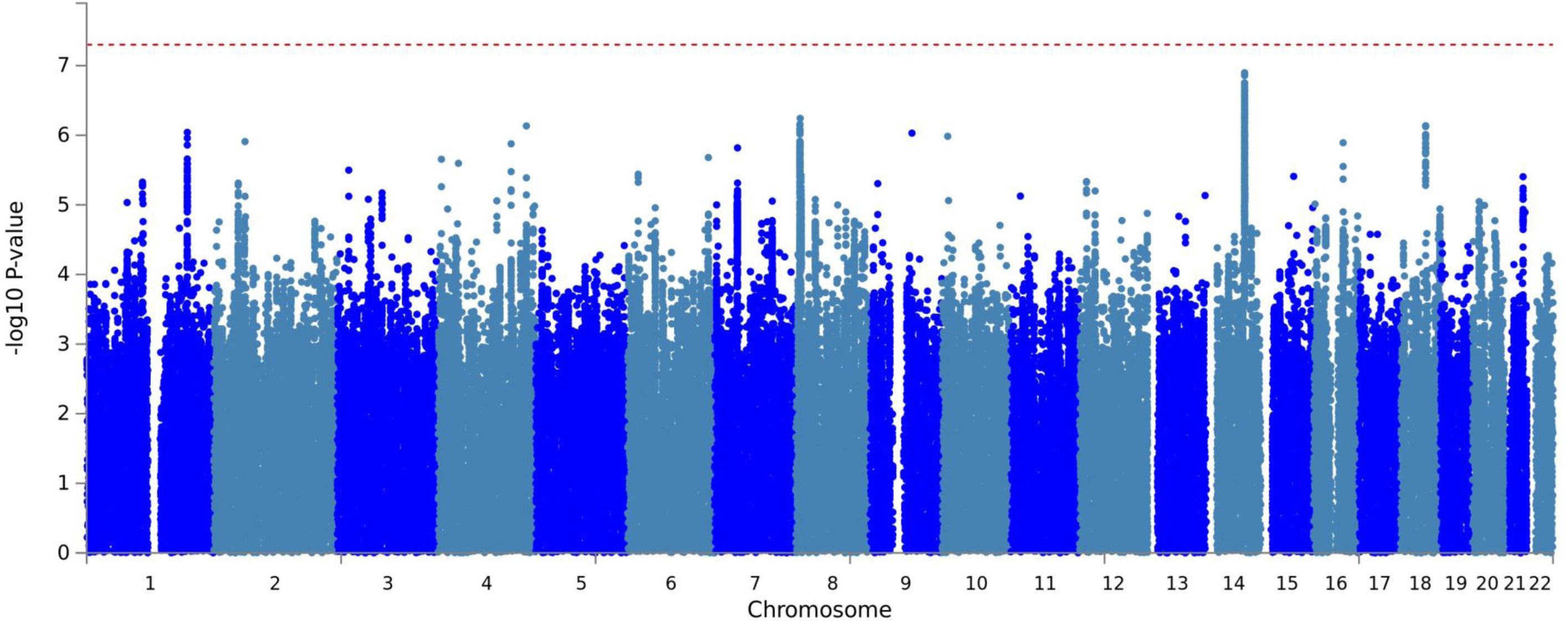
Manhattan plot of GWAS of mood instability in UK Biobank (females only).

**Figure S3.**
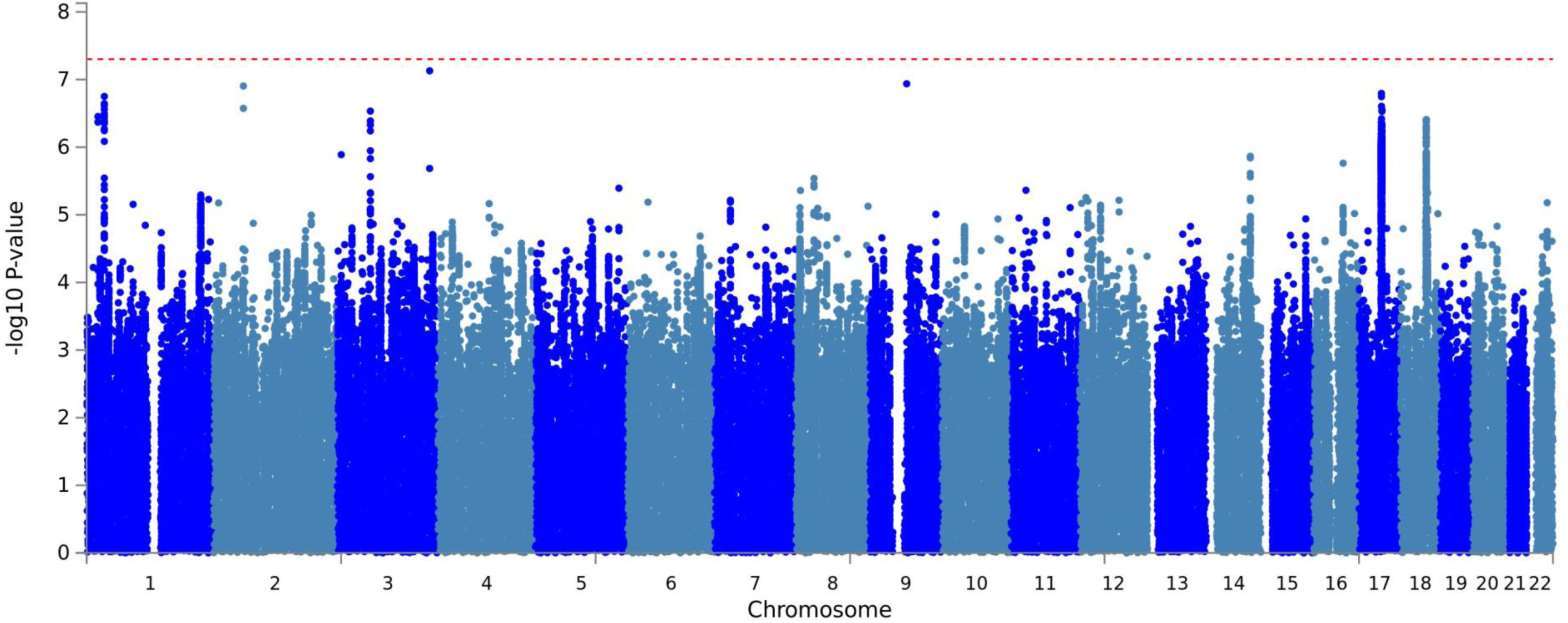
Manhattan plot of GWAS of mood instability in UK Biobank (age 58 and below).

**Figure S4.**
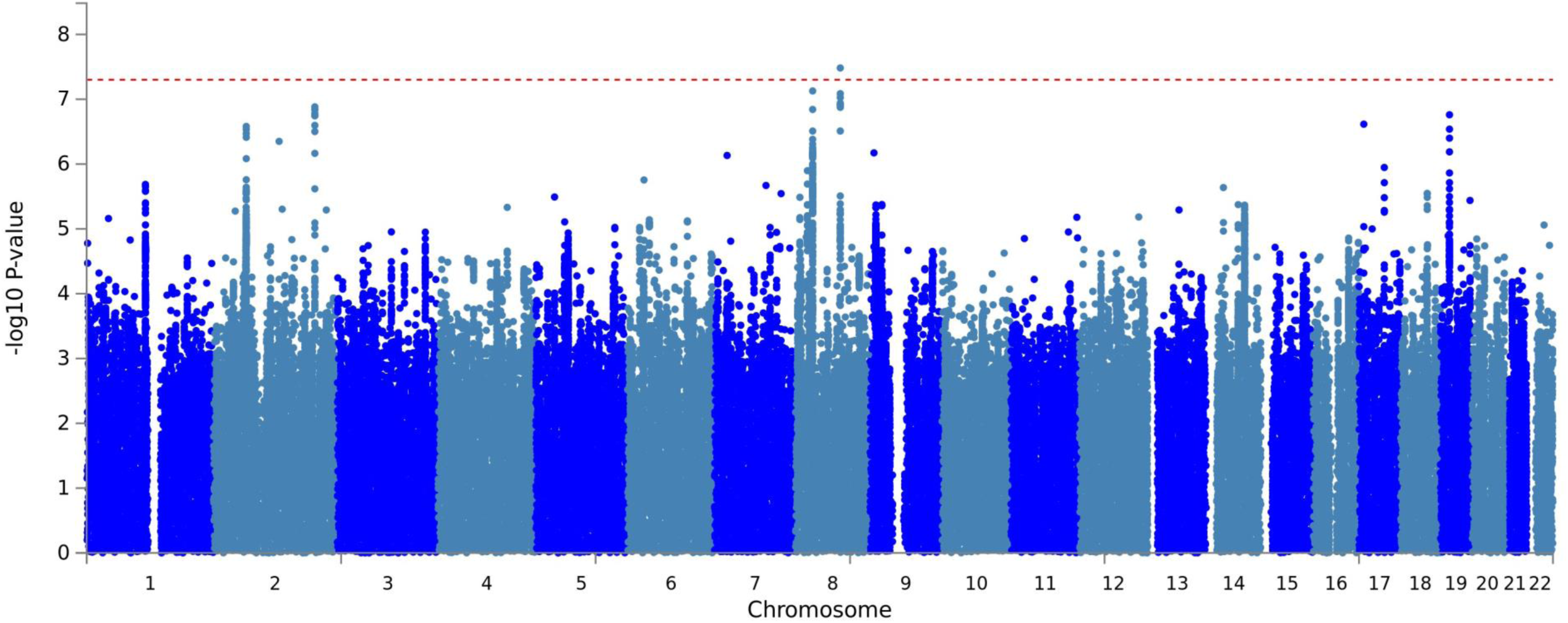
Manhattan plot of GWAS of mood instability in UK Biobank (age 59 and above).

## Supplementary material

**Table S1.** Genome-wide significant loci associated with mood instability in UK Biobank (excluding 9,865 participants with psychiatric disorder)

| Index SNP | Chr | Position | Risk Allele/Other Allele | RAF | Beta (SE) | P-value |
| --- | --- | --- | --- | --- | --- | --- |
| rs7829975 | 8 | 8,548,117 | A/T | 0.523 | 0.052 (0.009) | $5.32 \times 10^{-9}$ |
| rs10959826 | 9 | 11,459,410 | G/A | 0.789 | 0.055 (0.01) | $4.77 \times 10^{-7}$ |
| rs397852991 | 14 | 75,268,920 | C/CA | 0.673 | 0.045 (0.009) | $1.25 \times 10^{-6}$ |
| rs8084280 | 18 | 50,726,749 | T/A | 0.514 | 0.047 (0.008) | $1.35 \times 10^{-7}$ |

*Shown are LD-independent genome-wide significant SNP associations for mood instability (sorted by genomic position according to NCBI Build 37). Chromosome (Chr) and Position denote the location of the index SNP. RAF = risk allele frequency. Beta = logistic regression coefficient for allele1, SE = standard error for Beta. P-value = the probability of getting the derived test statistic under the null hypothesis. The final column indicates protein-coding genes at the associated loci (see regional plots in supplementary information) or, where there are no genes at the associated locus, the nearest gene if less than 1 MB from the locus.*

**Table S2.** Psychiatric polygenic risk score analysis of mood instability (adjusted for age, sex, genotyping chip and PGCs 1-8; n_total_=104,103, n_cas_=43,660, n_con_=60,443)

| Predictor | P | Beta | SE | OR | conf lower | conf upper | Nagelkerke r<sup>2</sup> |
| --- | --- | --- | --- | --- | --- | --- | --- |
| MDD_0.01 | 1.88*10<sup>-11</sup> | 0.0148 | 0.00221 | 1.0149 | 1.011 | 1.0193 | 0.0216 |
| MDD_0.05 | 5.47*10<sup>-22</sup> | 0.0216 | 0.00224 | 1.0218 | 1.02 | 1.0263 | 0.0221 |
| MDD_0.1 | 1.11*10<sup>-26</sup> | 0.0242 | 0.00226 | 1.0244 | 1.02 | 1.029 | 0.0224 |
| MDD_0.5 | 1.00*10<sup>-34</sup> | 0.0281 | 0.00229 | 1.0285 | 1.02 | 1.0332 | 0.0229 |
| bipolar_gws | 3.64*10<sup>-01</sup> | 0.00201 | 0.00221 | 1.002 | 0.99768 | 1.0064 | 0.021 |
| bipolar_0.01 | 1.67*10<sup>-01</sup> | 0.00305 | 0.00221 | 1.0031 | 0.99873 | 1.0074 | 0.021 |
| bipolar_0.05 | 1.28*10<sup>-01</sup> | 0.00338 | 0.00222 | 1.0034 | 0.99903 | 1.0078 | 0.021 |
| bipolar_0.1 | 1.65*10<sup>-01</sup> | 0.00311 | 0.00224 | 1.0031 | 0.99872 | 1.0075 | 0.021 |
| bipolar_0.5 | 1.00*10<sup>-01</sup> | 0.0037 | 0.00225 | 1.0037 | 0.99929 | 1.0081 | 0.021 |
| SCZ_gws | 1.08*10<sup>-01</sup> | 0.0035 | 0.00218 | 1.0035 | 0.99924 | 1.0078 | 0.021 |
| SCZ_0.01 | 1.55*10<sup>-03</sup> | 0.00718 | 0.00227 | 1.0072 | 1.0027 | 1.0117 | 0.0211 |
| SCZ_0.05 | 1.19*10<sup>-04</sup> | 0.00885 | 0.0023 | 1.0089 | 1.0044 | 1.0134 | 0.0211 |
| SCZ_0.1 | 6.71*10<sup>-05</sup> | 0.0092 | 0.00231 | 1.0092 | 1.0047 | 1.0138 | 0.0212 |
| SCZ_0.5 | 1.24*10<sup>-04</sup> | 0.00893 | 0.00233 | 1.009 | 1.0044 | 1.0136 | 0.0212 |

Shown are the results of a logistic regression using psychiatric PRS over a range of P value cut offs split into deciles. Predictor = The PRS used as a predictor in the model in the format "Psychiatric condition \_ p value cut off", P= the P value of the PRS predictor, Beta= the coefficient of the PRS predictor, SE = the standard error of the PRS predictor, OR= the odds ratio of the PRS predictor, conf lower and conf upper = the upper and lower 95% confidence interval of the PRS predictor, Nagelkerke r<sup>2</sup> = the variance explained by the whole model.

**Table S3.** Psychiatric polygenic risk score analysis of mood instability in females (adjusted for age, genotyping chip and PGCs 1-8; n_total_=53,279, n_cas_=23,308, n_con=_ 29,971)

| Predictor | P | Beta | SE | OR | conf lower | conf upper | Nagelkerke R2 |
| --- | --- | --- | --- | --- | --- | --- | --- |
| MDD_0.01 | 8.80*10<sup>-06</sup> | 0.0137 | 0.00308 | 1.0138 | 1.0077 | 1.0199 | 0.024668 |
| MDD_0.05 | 1.17*10<sup>-10</sup> | 0.0201 | 0.00312 | 1.0203 | 1.0141 | 1.0266 | 0.025205 |
| MDD_0.1 | 2.96*10<sup>-12</sup> | 0.022 | 0.00315 | 1.0222 | 1.0159 | 1.0286 | 0.025383 |
| MDD_0.5 | 2.89*10<sup>-17</sup> | 0.027 | 0.0032 | 1.0274 | 1.021 | 1.0338 | 0.025945 |
| bipolar_gws | 6.20*10<sup>-01</sup> | 0.00153 | 0.00309 | 1.0015 | 0.99549 | 1.0076 | 0.024186 |
| bipolar_0.01 | 5.35*10<sup>-02</sup> | 0.00593 | 0.00307 | 1.0059 | 0.99991 | 1.012 | 0.024272 |
| bipolar_0.05 | 1.01*10<sup>-01</sup> | 0.00509 | 0.0031 | 1.0051 | 0.99901 | 1.0112 | 0.024246 |
| bipolar_0.1 | 1.27*10<sup>-01</sup> | 0.00475 | 0.00312 | 1.0048 | 0.99864 | 1.0109 | 0.024237 |
| bipolar_0.5 | 6.92*10<sup>-02</sup> | 0.00569 | 0.00313 | 1.0057 | 0.99955 | 1.0119 | 0.024261 |
| SCZ_gws | 3.43*10<sup>-01</sup> | 0.00287 | 0.00303 | 1.0029 | 0.99694 | 1.0088 | 0.024202 |
| SCZ_0.01 | 1.62*10<sup>-02</sup> | 0.00759 | 0.00316 | 1.0076 | 1.0014 | 1.0139 | 0.024323 |
| SCZ_0.05 | 7.32*10<sup>-03</sup> | 0.00857 | 0.0032 | 1.0086 | 1.0023 | 1.015 | 0.024357 |
| SCZ_0.1 | 5.90*10<sup>-03</sup> | 0.00885 | 0.00321 | 1.0089 | 1.0026 | 1.0153 | 0.024367 |
| SCZ_0.5 | 1.47*10<sup>-02</sup> | 0.00791 | 0.00324 | 1.0079 | 1.0016 | 1.0144 | 0.024327 |

Shown are the results of a logistic regression using psychiatric PRS over a range of P value cut offs split into deciles. Predictor = The PRS used as a predictor in the model in the format "Psychiatric condition \_ p value cut off", P= the P value of the PRS predictor, Beta= the coefficient of the PRS predictor, SE = the standard error of the PRS predictor, OR= the odds ratio of the PRS predictor, conf lower and conf upper = the upper and lower 95% confidence interval of the PRS predictor, Nagelkerke r2 = the variance explained by the whole model.

**Table S4.** Psychiatric polygenic risk score analysis of mood instability in males only (adjusted for age, genotyping chip and PGCs 1-8, n_total_=50,824, n_cas_=24,804, n_con_=27,939)

| Predictor | P | Beta | SE | OR | conf lower | conf upper | Nagelkerke R2 |
| --- | --- | --- | --- | --- | --- | --- | --- |
| MDD_0.01 | 4.42*10<sup>-07</sup> | 0.016 | 0.00317 | 1.0162 | 1.0099 | 1.0225 | 0.01567 |
| MDD_0.05 | 4.99*10<sup>-13</sup> | 0.023 | 0.00322 | 1.0235 | 1.0171 | 1.03 | 0.016372 |
| MDD_0.1 | 2.69*10<sup>-16</sup> | 0.027 | 0.00324 | 1.0269 | 1.0204 | 1.0334 | 0.016762 |
| MDD_0.5 | 3.01*10<sup>-19</sup> | 0.03 | 0.00329 | 1.0299 | 1.0233 | 1.0366 | 0.017115 |
| bipolar_gws | 4.30*10<sup>-01</sup> | 0.003 | 0.00317 | 1.0025 | 0.99629 | 1.0088 | 0.015015 |
| bipolar_0.01 | 9.96*10<sup>-01</sup> | -0.00001 | 0.00317 | 0.99999 | 0.99379 | 1.0062 | 0.014999 |
| bipolar_0.05 | 6.48*10<sup>-01</sup> | 0.001 | 0.00319 | 1.0015 | 0.99521 | 1.0077 | 0.015004 |
| bipolar_0.1 | 6.98*10<sup>-01</sup> | 0.001 | 0.00321 | 1.0012 | 0.99496 | 1.0076 | 0.015003 |
| bipolar_0.5 | 6.45*10<sup>-01</sup> | 0.001 | 0.00323 | 1.0015 | 0.99517 | 1.0079 | 0.015004 |
| SCZ_gws | 1.82*10<sup>-01</sup> | 0.004 | 0.00314 | 1.0042 | 0.99805 | 1.0104 | 0.015046 |
| SCZ_0.01 | 4.17*10<sup>-02</sup> | 0.007 | 0.00326 | 1.0067 | 1.0003 | 1.0131 | 0.015108 |
| SCZ_0.05 | 6.18*10<sup>-03</sup> | 0.009 | 0.00331 | 1.0091 | 1.0026 | 1.0157 | 0.015196 |
| SCZ_0.1 | 4.26*10<sup>-03</sup> | 0.01 | 0.00332 | 1.0095 | 1.003 | 1.0161 | 0.015214 |
| SCZ_0.5 | 2.80*10<sup>-03</sup> | 0.01 | 0.00335 | 1.0101 | 1.0035 | 1.0167 | 0.015234 |

Shown are the results of a logistic regression using psychiatric PRS over a range of P value cut offs split into deciles. Predictor = The PRS used as a predictor in the model in the format "Psychiatric condition \_ p value cut off", P= the P value of the PRS predictor, Beta= the coefficient of the PRS predictor, SE = the standard error of the PRS predictor, OR= the odds ratio of the PRS predictor, conf lower and conf upper = the upper and lower 95% confidence interval of the PRS predictor, Nagelkerke r2 = the variance explained by the whole model.

**Table S5.** Psychiatric polygenic risk score analysis of mood instability in those equal to or younger than the median age of 58 (adjusted for age, sex genotyping chip and PGCs 1-8; n_total_=52,743, n_cas_=24,804, n_con_=27,939)

| Predictor | P | Beta | SE | OR | conf lower | conf upper | Nagelkerke R2 |
| --- | --- | --- | --- | --- | --- | --- | --- |
| MDD_0.01 | 1.80*10<sup>-06</sup> | 0.0146 | 0.00305 | 1.0147 | 1.0086 | 1.0208 | 0.012342 |
| MDD_0.05 | 9.65*10<sup>-13</sup> | 0.0221 | 0.0031 | 1.0224 | 1.0162 | 1.0286 | 0.013048 |
| MDD_0.1 | 8.28*10<sup>-16</sup> | 0.0252 | 0.00313 | 1.0255 | 1.0192 | 1.0318 | 0.013397 |
| MDD_0.5 | 1.13*10<sup>-22</sup> | 0.0311 | 0.00317 | 1.0316 | 1.0252 | 1.038 | 0.014181 |
| bipolar_gws | 8.77*10<sup>-01</sup> | 0.000475 | 0.00306 | 1.0005 | 0.99449 | 1.0065 | 0.011771 |
| bipolar_0.01 | 2.51*10<sup>-01</sup> | 0.00351 | 0.00305 | 1.0035 | 0.99753 | 1.0095 | 0.011803 |
| bipolar_0.05 | 1.35*10<sup>-01</sup> | 0.00459 | 0.00308 | 1.0046 | 0.99857 | 1.0107 | 0.011826 |
| bipolar_0.1 | 2.66*10<sup>-01</sup> | 0.00344 | 0.00309 | 1.0034 | 0.99738 | 1.0095 | 0.011801 |
| bipolar_0.5 | 1.44*10<sup>-01</sup> | 0.00454 | 0.00311 | 1.0046 | 0.99844 | 1.0107 | 0.011824 |
| SCZ_gws | 1.73*10<sup>-02</sup> | 0.00719 | 0.00302 | 1.0072 | 1.0013 | 1.0132 | 0.011912 |
| SCZ_0.01 | 7.10*10<sup>-03</sup> | 0.00845 | 0.00314 | 1.0085 | 1.0023 | 1.0147 | 0.011952 |
| SCZ_0.05 | 4.49*10<sup>-04</sup> | 0.0112 | 0.00318 | 1.0112 | 1.0049 | 1.0176 | 0.012079 |
| SCZ_0.1 | 4.34*10<sup>-04</sup> | 0.0112 | 0.0032 | 1.0113 | 1.005 | 1.0177 | 0.012081 |
| SCZ_0.5 | 5.86*10<sup>-04</sup> | 0.0111 | 0.00323 | 1.0112 | 1.0048 | 1.0176 | 0.012067 |

Shown are the results of a logistic regression using psychiatric PRS over a range of P value cut offs split into deciles. Predictor = The PRS used as a predictor in the model in the format "Psychiatric condition \_ p value cut off", P= the P value of the PRS predictor, Beta= the coefficient of the PRS predictor, SE = the standard error of the PRS predictor, OR= the odds ratio of the PRS predictor, conf lower and conf upper = the upper and lower 95% confidence interval of the PRS predictor, Nagelkerke r2 = the variance explained by the whole model.

**Table S6.**
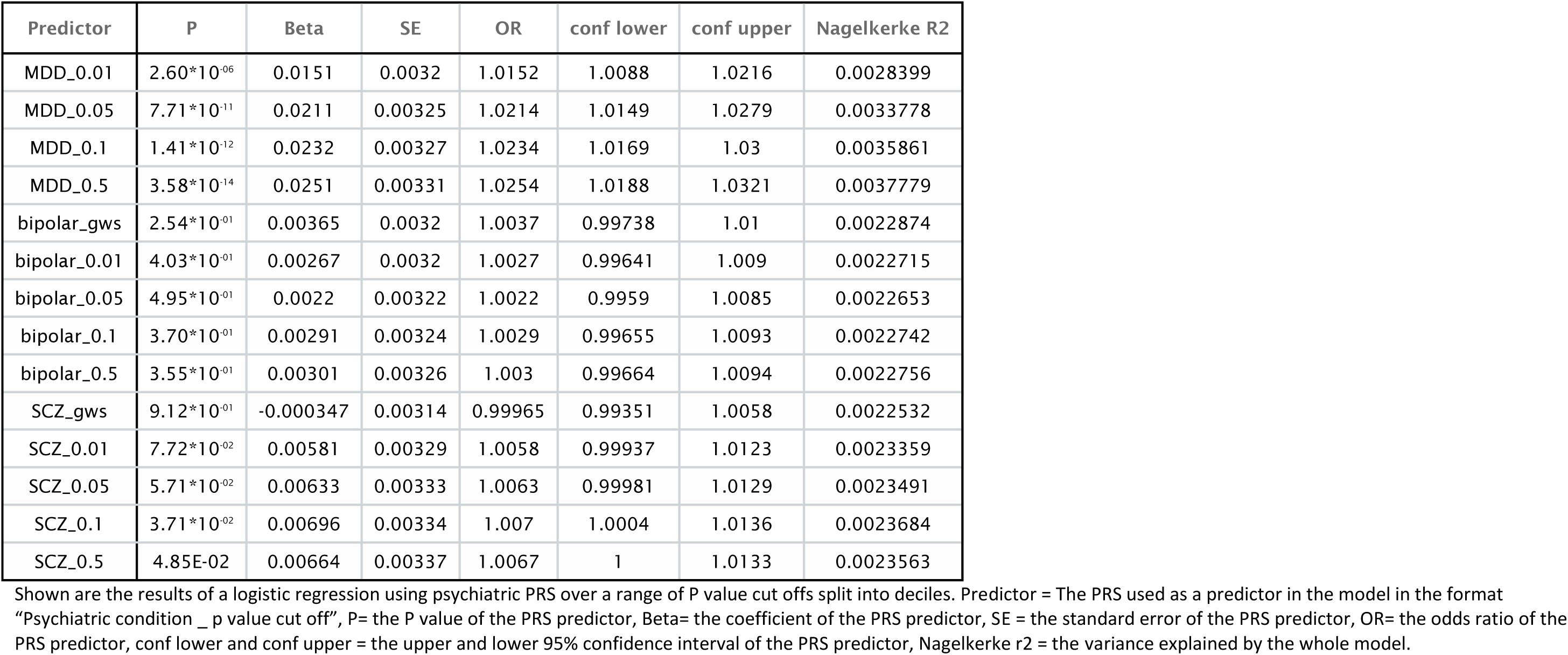
Psychiatric polygenic risk score analysis of mood instability in those older than the median age of 58 (adjusted for age, sex genotyping chip and PGCs 1-8, n_total_=51,360, n_cas_=18,856, n_con_=23,504)

| Predictor | P | Beta | SE | OR | conf lower | conf upper | Nagelkerke R2 |
| --- | --- | --- | --- | --- | --- | --- | --- |
| MDD_0.01 | $2.60 \times 10^{-6}$ | 0.0151 | 0.0032 | 1.0152 | 1.0088 | 1.0216 | 0.0028399 |
| MDD_0.05 | $7.71 \times 10^{-11}$ | 0.0211 | 0.00325 | 1.0214 | 1.0149 | 1.0279 | 0.0033778 |
| MDD_0.1 | $1.41 \times 10^{-12}$ | 0.0232 | 0.00327 | 1.0234 | 1.0169 | 1.03 | 0.0035861 |
| MDD_0.5 | $3.58 \times 10^{-14}$ | 0.0251 | 0.00331 | 1.0254 | 1.0188 | 1.0321 | 0.0037779 |
| bipolar_gws | $2.54 \times 10^{-01}$ | 0.00365 | 0.0032 | 1.0037 | 0.99738 | 1.01 | 0.0022874 |
| bipolar_0.01 | $4.03 \times 10^{-01}$ | 0.00267 | 0.0032 | 1.0027 | 0.99641 | 1.009 | 0.0022715 |
| bipolar_0.05 | $4.95 \times 10^{-01}$ | 0.0022 | 0.00322 | 1.0022 | 0.9959 | 1.0085 | 0.0022653 |
| bipolar_0.1 | $3.70 \times 10^{-01}$ | 0.00291 | 0.00324 | 1.0029 | 0.99655 | 1.0093 | 0.0022742 |
| bipolar_0.5 | $3.55 \times 10^{-01}$ | 0.00301 | 0.00326 | 1.003 | 0.99664 | 1.0094 | 0.0022756 |
| SCZ_gws | $9.12 \times 10^{-01}$ | -0.000347 | 0.00314 | 0.99965 | 0.99351 | 1.0058 | 0.0022532 |
| SCZ_0.01 | $7.72 \times 10^{-02}$ | 0.00581 | 0.00329 | 1.0058 | 0.99937 | 1.0123 | 0.0023359 |
| SCZ_0.05 | $5.71 \times 10^{-02}$ | 0.00633 | 0.00333 | 1.0063 | 0.99981 | 1.0129 | 0.0023491 |
| SCZ_0.1 | $3.71 \times 10^{-02}$ | 0.00696 | 0.00334 | 1.007 | 1.0004 | 1.0136 | 0.0023684 |
| SCZ_0.5 | 4.85E-02 | 0.00664 | 0.00337 | 1.0067 | 1 | 1.0133 | 0.0023563 |

Shown are the results of a logistic regression using psychiatric PRS over a range of P value cut offs split into deciles. Predictor = The PRS used as a predictor in the model in the format "Psychiatric condition \_ p value cut off", P= the P value of the PRS predictor, Beta= the coefficient of the PRS predictor, SE = the standard error of the PRS predictor, OR= the odds ratio of the PRS predictor, conf lower and conf upper = the upper and lower 95% confidence interval of the PRS predictor, Nagelkerke r2 = the variance explained by the whole model.

